# Surveying armadillo and bat trypanosomes by DNA metabarcoding with Oxford Nanopore Technologies sequencing: the importance of fine-tuning parameters to identify mixed infections

**DOI:** 10.64898/2026.08.03.742417

**Authors:** C. Miguel Pinto, Manuel Calvopiña, Sofía Ocaña-Mayorga, Daniel Romero-Alvarez, Carlos Bastidas-Caldes, Pamela Lojan-Cueva, Daniela Reyes-Barriga D, Alejandra Bedoya-Jaramillo, Víctor Romero, Nicté Ordoñez-Garza, Andrea Au-Hing A, Mónica Páez-Vacas, Julio Carrión-Olmedo, Ricardo S. P. Patiño, Pablo Jarrín-V

## Abstract

**Background:** The ecological dynamics between *Trypanosoma* parasites and their wild mammalian hosts, such as bats and armadillos, are complex. Recent 18S rRNA metabarcoding studies have reported extraordinary levels of hidden parasite diversity and frequent multi-lineage coinfections within individual wild hosts. However, the boundary between genuine biological coinfection and methodological artifact remains difficult to establish. Based on Gause’s principle of competitive exclusion, the mammalian bloodstream represents a highly constrained niche where stable coexistence of identical ecological competitors is theoretically rare. We hypothesize that previously reported hyper-diverse *Trypanosoma* coinfections are largely bioinformatic artifacts, and that true intra-host dynamics instead favor single-lineage dominance.

**Methods:** To test this hypothesis, we sequenced samples from 27 wild armadillos (*Dasypus novemcinctus*) and 26 bats from Ecuador. The 18S rRNA gene was amplified via nested PCR and sequenced using an Oxford Nanopore Technologies MinION platform. We developed a progressively stringent bioinformatics pipeline to evaluate coinfection hypotheses. Raw reads were processed through three alignment scenarios: Lenient, Moderate, and Conservative. These scenarios modulate sequence identity, mapping quality (MAPQ), and coverage thresholds to effectively isolate true biological signals from alignment ambiguity.

**Results:** Under lenient alignment parameters, the resulting profiles mirrored previous literature, exhibiting massive apparent intra-host multi-lineage diversity. However, as bioinformatic stringency increased to conservative thresholds (≥ 98% sequence identity, ≥ 99% coverage, and MAPQ ≥ 30), artifactual pseudo-coinfections collapsed. The highly restricted dataset demonstrated overwhelming single-lineage dominance, validating only three active mixed infections out of the retained samples. Furthermore, our rigorous pipeline isolated rare but genuine biological signals, including the detection of *Trypanosoma cruzi marinkellei*—historically considered a bat-restricted subgenus—within the terrestrial armadillo cohort. We also confirmed the presence of *T. cruzi* DTU III (TcIII) in Ecuadorian armadillos, representing a significant biogeographical record for the region.

**Conclusions:** Once methodological noise is computationally stripped away, active multi-strain *Trypanosoma coinfections* in the host bloodstream are revealed to be ecologically anomalous. Our findings strongly support the principle of competitive exclusion, suggesting established lineages actively suppress competitors. While Oxford Nanopore sequencing offers necessary resolution for wildlife parasitology, fine-tuning algorithmic parameters is critical to accurately represent host-parasite networks and prevent the artificial inflation of intra-host diversity metrics.

**Author summary:** Previous studies using DNA metabarcoding have reported that wild mammals, such as bats, frequently harbor complex communities of multiple *Trypanosoma* parasite lineages simultaneously. However, ecological principles suggest that identical competitors struggle to coexist stably within a constrained environment like the host bloodstream. To investigate whether these reported high coinfection rates reflect true biology or methodological artifacts, we sequenced the 18S rRNA gene of *Trypanosoma* from 26 bats and 27 armadillos in Ecuador. We processed the sequencing data through computational pipelines with progressively stricter filtering parameters. We observed that under lenient filtering, animals appeared to have highly diverse, mixed infections. Conversely, when strict parameters were applied to remove potential analytical noise, the artificial complexity collapsed, revealing that the vast majority of hosts were dominated by a single parasite lineage. We confirmed only three active mixed infections in our highly restricted dataset. Our findings indicate that active multi-strain *Trypanosoma* coinfections are rare, aligning with the principle of competitive exclusion. These results highlight the necessity of applying rigorous bioinformatic filters to accurately evaluate host-parasite interactions and avoid overestimating diversity metrics.

## Introduction

*Trypanosoma* parasites are a fairly diverse group of blood parasites of all classes of vertebrates [1]. The complex ecological dynamics between vertebrate hosts, their arthropod vectors, and these hemoflagellate parasites have driven remarkable host-parasite associations. For example, bats serve as ancient and highly diverse hosts for numerous species and genotypes within the genus *Trypanosoma*, particularly within the speciose *T. cruzi* clade [2,3]. On the other hand, armadillos (order Cingulata) are considered important hosts of certain lineages of the medically important *Trypanosoma cruzi* [4]. Armadillos act as wild reservoirs for T. cruzi DTU TcIII, maintaining transmission cycles that are largely isolated from domestic environments [5–7]. Notably, infection rates in these populations can be high, reaching 57.7% in *Dasypus novemcinctus* [7] and 65–76% in *Zaedyus pichiy* [8,9].

Historically, uncovering the diversity patterns of infection have mostly relied on standard analysis of discrete genetic markers, often the genes 18S rRNA and GAPDH [10–14]. More recently, 18S rRNA metabarcoding has revealed extraordinary levels of hidden trypanosome diversity and mixed infections circulating within wild bat populations [15,16]. Consequently, a prevailing narrative has emerged in the literature suggesting that multiple *Trypanosoma* lineages frequently co-inhabit individual bat hosts [10,14–16]. However, characterizing these previous findings as definitive evidence of widespread coinfection may overstate the biological reality [16–20].

While these foundational studies highlighted broad trends regarding *Trypanosoma* genetic signatures, establishing the boundary between a true biological coinfection and a methodological artifact remains challenging [17]. Issues such as PCR amplification biases, chimeric sequences, sequence references and sequence alignments, and the complex systematic concepts surrounding hybrid populations [17] can artificially inflate estimates of intra-host diversity [20]. This concern is not exclusive to bat studies. In non-volant sylvatic mammals, field surveys consistently report single-lineage dominance such as in armadillos that have been reported predominantly infected by TcIII and opossums by TcI [5–7]. This host specific pattern suggests a limited multi-lineage coexistence across different host groups.

Recent exploratory studies emphasize the utility of Neotropical bats as sentinels for emerging zoonoses, including *Trypanosoma cruzi*, utilizing broad-spectrum metagenomic next-generation sequencing (mNGS) [21]. However, the untargeted nature of such sequencing can inflate the risk of false positives, often necessitating secondary confirmation via targeted tools like qPCR to validate active pathogen presence [21]. While these foundational studies highlighted broad trends regarding *Trypanosoma* genetic signatures, establishing the boundary between a true biological coinfection and a methodological artifact remains challenging. Issues such as PCR amplification biases, chimeric sequences, and alignment ambiguity can artificially inflate estimates of intra-host diversity [21].

From an ecological perspective, the mammalian bloodstream should represent a highly constrained and competitive niche [22]. According to Gause’s principle of competitive exclusion, pathogen species sharing identical ecological niches cannot coexist stably and indefinitely; the more competitive species will eventually take over and exclude the other [23]. Within a host, this exclusion arises through asymmetric competition, primarily driven by exploitative competition for shared host energy resources or apparent competition mediated by shared immune cells [24]. For coinfection to successfully occur, each parasite species must exert a stronger negative effect on its own growth rate (intraspecific competition) than on its competitor’s (interspecific competition), a balance rarely achieved in such restricted environments [24]. Furthermore, evolutionary instability often arises due to conflicts of interest between coinfecting parasites, pushing the system toward exclusion rather than harmony [25–28].

Experimental evidence strongly supports these theoretical constraints. In analogous blood-parasite systems like *Plasmodium*, mixed infections of different clones result in the dominant clone rapidly outgrowing and competitively suppressing the weaker clone, sometimes completely excluding it to undetectable levels within days [29]. This pattern holds across micro-ecological systems, where competitive exclusion is the overwhelming outcome of multi-parasite inoculations, and actual cellular coinfection remains an extreme rarity, occurring in less than 1% of infected cells [30]. Recent evidence suggests that biological coinfections in neotropical wildlife remain infrequent and are restricted to highly specific lineage pairings in bats [5–7,20]. Therefore, we hypothesize that true, active *Trypanosoma* coinfections in bat blood are ecologically and evolutionarily anomalous. We propose that the high coinfection rates previously reported [16,17,19] are largely bioinformatic artifacts of lower-resolution sequencing pipelines, or represent transient, unstable encounters rather than established multi-strain communities.

In this study, we compare the tuning parameters, from generalistic to highly restrictive, in a nanopore-based bioinformatics pipeline to assess the prevalence of *Trypanosoma* coinfections in blood samples of armadillos and bats. By filtering for high-confidence, full-length alignments, we aim to demonstrate that once methodological noise is stripped away, the resulting infection [31] profiles are highly consistent with the ecological expectations of competitive exclusion, presenting a strong trend toward single-lineage dominance rather than widespread coinfection.

## 2. Methods

### 2.1. Host Sampling

Armadillo Sampling: Tissue samples (spleen, liver, or muscle) were collected from the bushmeat catches of local hunters between 2021 and 2022. Three additional samples were available from the collection of the Instituto Nacional de Biodiversidad in Ecuador (INABIO). Tissues were stored in 70% ethanol. These samples were previously analyzed for the molecular diagnosis of *Mycobacterium leprae* and *M. lepromatosis* [31]. Access to these samples was approved by the Ministerio del Ambiente, Agua y Transición Ecológica under the access contract to genetic resources MAATE-DBI-CM-2023-0334. All samples were taxonomically assigned to *Dasypus novemcinctus*, as sequences shared 100% identity with a reference genome [31].

Bat Sampling: Bats were sampled in the Cuyabeno Wildlife Reserve, Ecuadorian Amazon, during the 2019 rainy season. Between June 1st and 4th, six mist nets (3 × 9 m) were deployed along forest edges, trails, and water bodies. A total of 107 bats were captured, of which 26 samples were sequenced. From each bat, a small amount of blood was drawn from the antebrachial vein, using a 26 gauge needle [32]. Once the vein was punctured, the blood was collected on an FTA card strip, and after air drying it was stored inside a cryovial tube. Bat taxonomic identification was based on external morphological and morphometric characters following [33]. After sampling and identification, bats were released at the capture site. Capture and handling procedures followed the guidelines of the American Society of Mammalogists [34] and had the Ministry of Environment authorization MAE-DNB-CM-2017-0068.

### 2.2. DNA extraction and PCR amplifications

DNA from armadillo tissues was extracted in January 2024 using the Wizard Genomic DNA Purification Kit (Promega), following the manufacturer’s protocol for animal tissue. DNA from bat blood samples was extracted in March 2024 using the PureLink Genomic DNA Mini Kit (Invitrogen), following the manufacturer’s protocol. For both sample types, DNA quality and concentration were assessed by NanoDrop spectrophotometry and extracted DNA was stored at -20°C until analysis.

To amplify the 18S rRNA gene, we followed a nested PCR approach [35] with a modification to the original protocol. The initial PCR amplification was performed using the primers SSU4_F (GTGCCAGCACCCGCGGTAAT) and 18Sq1R (CCACCGACCAAAAGCGGCCA) [11]; whereas the second PCR amplification employed the primers SSU561F and SSU561R. Both nested PCR amplifications were carried out with a touchdown PCR profile [36].

### 2.3. DNA sequencing

PCR amplification products were barcoded and multiplexed using the Rapid Barcoding Sequencing Kit 96 V14 (SQK-RBK114.96; Oxford Nanopore Technologies) under the manufacturer’s protocol. The multiplexed library was sequenced on a portable MinION Mk1C sequencing platform using R10.4.1 Flongle Flow Cells. The sequencing run was controlled and monitored using MinKNOW software (v24.11.8; MinKNOW Core v6.2.6) operating under the Bream protocol (v8.2.5). The experiment was configured for a 24-hour execution period, with automated pore scans programmed at regular 2-hour intervals to ensure optimal cell performance.

Raw ionic current signal data were stored in POD5 format, generating automated batches every hour or upon reaching 500,000,000 bases. Subsequent basecalling and demultiplexing were performed using Dorado (v7.6.7) with the High-accuracy model (v4.3.0) calibrated for a standard translocation speed of 400 bps. To ensure high data quality, a strict minimum quality filter was applied, discarding reads with a Q-score < 8. During demultiplexing, barcode trimming was enabled at the ends of valid sequences, while mid-read barcode filtering and modified basecalling were disabled. High-quality processed reads were exported into FASTQ format files at 10-minute intervals for downstream genomic analysis.

### 2.4. Creation of a reference database for the identification of *Trypanosoma* species based on the 18S rRNA region

To establish a robust and comprehensive reference framework for the accurate taxonomic assignment of *Trypanosoma* lineages, a curated reference database targeting the 18S ribosomal RNA (rRNA) gene was constructed. Initially, all available 18S rRNA sequences associated with the genus *Trypanosoma* were retrieved from the National Center for Biotechnology Information (NCBI) GenBank database on May 21st, 2025. This comprehensive retrieval generated an initial highly-redundant dataset comprising 2,097 sequences (trypanosoma_18S_db_2097.fas, S1 Data).

Given the high level of genetic redundancy inherent in public repositories, a dereplication step was essential to streamline downstream computational efficiency, mitigate the overrepresentation of clonal lineages, and ensure a balanced sequence-structure phylogeny. To achieve this, the initial FASTA dataset was processed utilizing the CD-HIT-EST sequence clustering algorithm [37]. A stringent sequence identity threshold of 99% (-c 0.99) was applied to cluster highly similar sequences and extract a single representative sequence per cluster. This rigorous clustering procedure effectively collapsed the initial dataset, yielding a curated, non-redundant reference database comprising 336 unique 18S rRNA *Trypanosoma* sequences (trypanosoma_18S_db_336.fas, S2 Data).

Then, we manually curated this dataset by interactions of alignments and removing redundant sequences (assumed to be the same species), with the exception of *Trypanosoma cruzi* were one sequence of each of the nine known main lineages was retained (i.e., DTUI to DTUVI, Tcbat, *T. cruzi marinkellei* I, and *T. cruzi marinkellei* II) [38,39]. We also trimmed the alignment at both extremes of the sequence to cover a fragment of ca. 560 bp (unaligned) that was amplified by the nested PCR protocol of [35,40]. The resulting final reference database consisted of 113 unique sequences. This optimized, dereplicated, and manually curated database served as the foundational reference architecture for all subsequent nanopore read mapping and taxonomic identification procedures (trypanosoma_18S_db_113.fas, S3 Data).

### 2.5. Bioinformatics Pipeline

#### 2.5.1. Quality Control and Sequence Filtering

Prior to alignment, raw Oxford Nanopore sequencing reads underwent stringent quality control and length filtering to minimize methodological noise and eliminate spurious sequences. Initial processing of the raw FASTQ files was performed using Chopper [41]. To ensure high confidence in base-calling accuracy, sequences with a mean Phred quality score below 10 (representing <90% expected base accuracy) were strictly discarded. Furthermore, Chopper [41] allowed us to isolate the specific ∼523 bp target amplicon and remove truncated fragments or artificially concatenated chimeric reads, a highly restrictive length filter was applied. Only reads measuring between 450 and 650 base pairs were retained for downstream analysis. This pre-processing step generated individual, high-quality FASTQ files (*_clean.fastq) for each sampled bat, establishing a robust, standardized baseline of amplicon sequences for the subsequent detection of *Trypanosoma* lineages (Fig. 1). The repository for the studied sequences is described in the Supporting information and data availability section.

**Figure 1.**
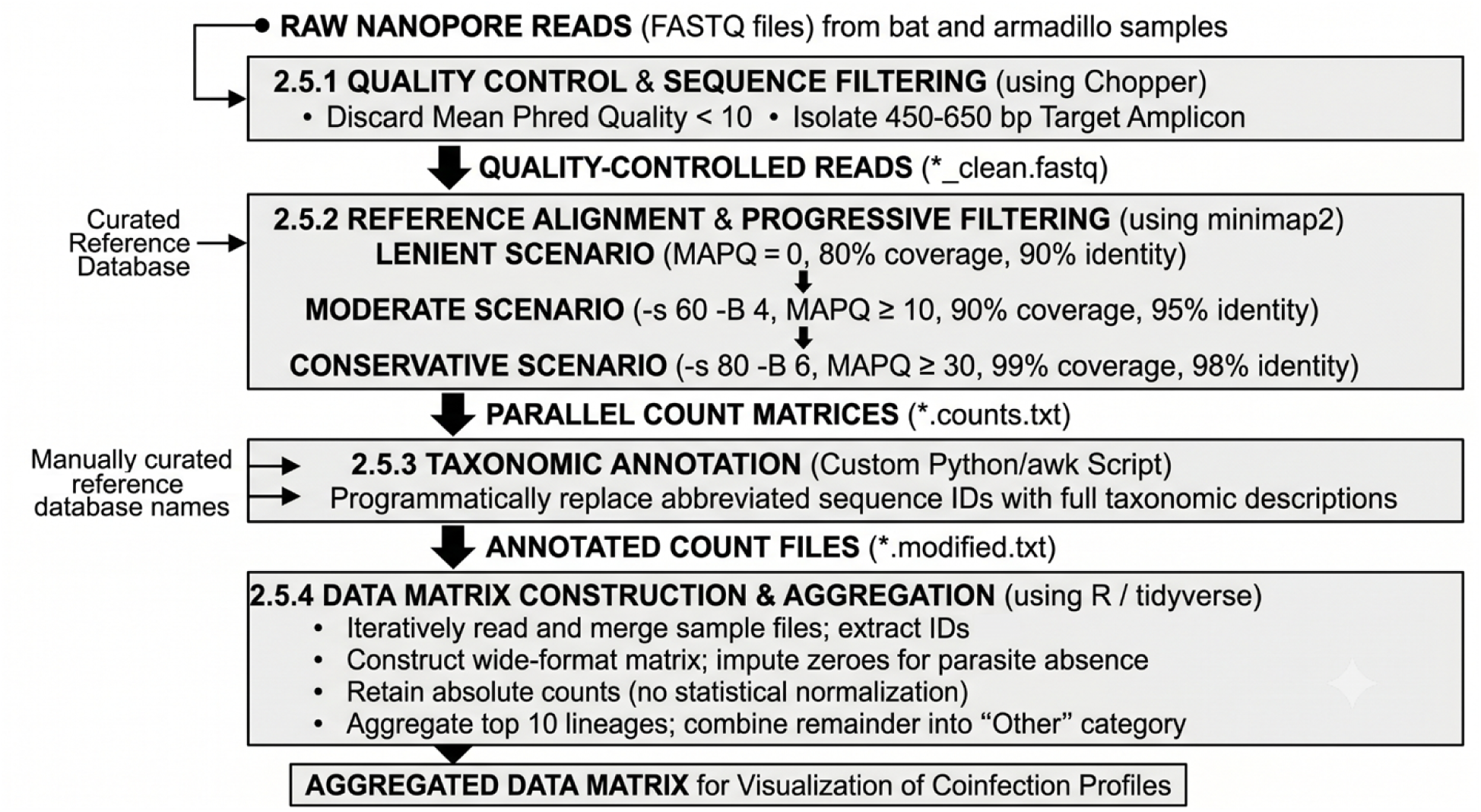
Workflow of the Oxford Nanopore 18S rRNA metabarcoding bioinformatics pipeline, with four primary analytical stages (named after each subsection in the manuscript): 1) initial read quality control and amplicon isolation, 2) reference alignment across three progressively restrictive filtering scenarios to evaluate coinfection hypotheses, 3) programmatic taxonomic annotation, and 4) data matrix aggregation in R using absolute read counts to visualize lineage dominance.

#### 2.5.2 Reference Alignment and Stringent Scenario-Based Filtering

To systematically test the hypothesis that previously reported high coinfection rates are largely bioinformatic artifacts driven by analytical noise, a comparative scenario analysis was conducted. We designed three distinct bioinformatic filtering pipelines, progressively modulating the stringency of the minimap2 [42] alignment algorithms and our custom sequence filters. This strategy was employed to emulate the spectrum of methodologies present in earlier literature and to isolate true biological signals from alignment ambiguity. The three generated scenarios were defined as follows (Fig. 2):

**Figure 2.**
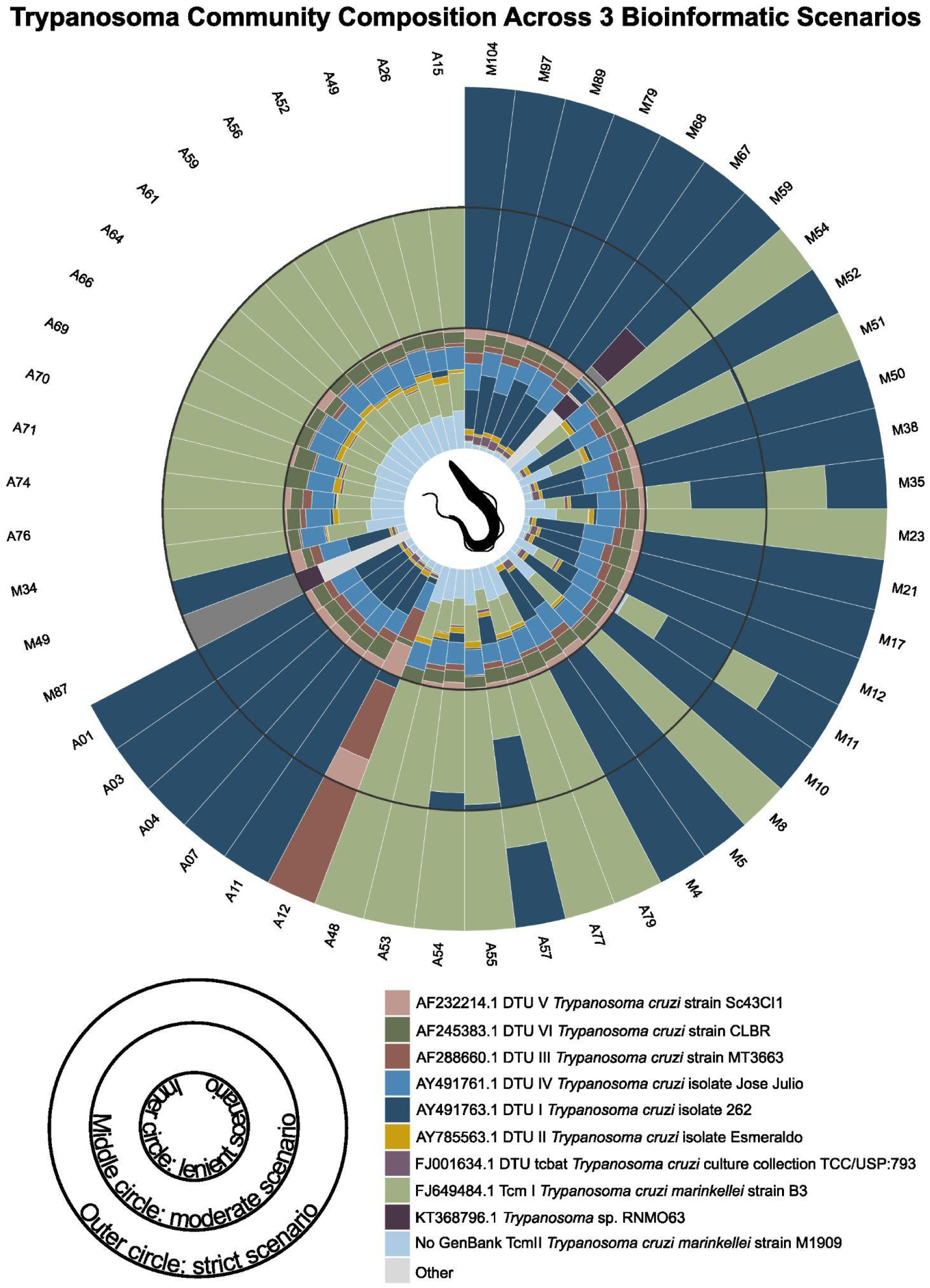
Hypotheses of *Trypanosoma* coinfection based on algorithmic filtering. Relative abundance of *Trypanosoma* strains per sample are displayed across three concentric rings, representing the relaxed (inner ring), moderate (middle ring), and conservative (outer ring) hypotheses of coinfection. Results are derived from the developed search, alignment, and filtering algorithm. Sample codes that start with A are derived from armadillos and those that start with M are samples from bats.

The Lenient Scenario (High-Noise Baseline): Designed to emulate lower-resolution methodologies and highly permissible clustering, this scenario maximized read retention by eliminating strict alignment penalties. The minimap2 algorithm was run using the default -cx map-ont preset without additional mismatch modifiers. During PAF filtering, the Mapping Quality (MAPQ) threshold was disabled (MAPQ = 0), allowing multi-mapping reads to be retained. The biological integrity thresholds were relaxed to accept reads with a minimum query or target coverage of 80% and a sequence identity of only 90%.

The Moderate Scenario (Standard Clustering): Designed to represent a standard environmental metabarcoding pipeline, this scenario applied intermediate stringency to clear fragmented reads while retaining standard taxonomic operational thresholds. Intermediate mismatch penalties were introduced to the alignment (-s 60 -B 4). The mapping confidence threshold was set to MAPQ ≥ 10 (approximately 90% confidence of unique mapping). The minimum coverage requirement was raised to 90%, and the sequence identity threshold was set to 95%, a standard cutoff often utilized for broad eukaryotic diversity assessments.

The Conservative Scenario (Highly Restrictive): Designed to test the competitive exclusion hypothesis, this baseline enforced absolute biological certainty to detect only true, stable lineage dominance. Stringent alignment modifiers were applied to heavily penalize base mismatches (-s 80 -B 6). The retention filter required an MAPQ ≥ 30, guaranteeing ∼99.9% confidence that the read mapped uniquely to a single specific reference sequence. Furthermore, an alignment was only counted if it achieved near-perfect sequence identity (≥ 98%; see S4 Text for a sensitivity analysis justifying this threshold) and covered at least 99% of either the query sequence or the target reference gene.

By iteratively processing the quality-controlled reads through these three structured paradigms, we generated parallel count matrices. The resulting variations in apparent multi-strain detection were subsequently evaluated to track the collapse of possibly artifactual pseudo-coinfections as methodological stringency increased (Fig. 1). The three versions of the pipeline are available at GitHub (See the supporting information and data availability section).

#### 2.5.3. Taxonomic Annotation and Data Formatting

Following the generation of individual sample count files, a custom Python-wrapped automated script (process_data.py) was utilized to annotate and format the output data. Because standard alignment outputs frequently truncate FASTA headers to their first discrete string, the initial count summaries contained abbreviated sequence identifiers. To prepare the data for biological interpretation, the script executed an awk-based dictionary lookup to restore the full taxonomic lineage descriptions. Specifically, the algorithm iterated through the filtered alignment summaries (*.counts.txt) provided by the previous minimap2 step and cross-referenced the abbreviated sequence identifiers against a manually curated reference database (reference_names.txt). Upon matching an identifier, the script programmatically replaced the abbreviated string with its corresponding full taxonomic and lineage descriptor, while preserving the associated sequence counts. This process yielded a new suite of fully annotated, tab-separated output files (*.modified.txt). This essential formatting step ensured that the high-confidence read counts for each individual bat sample were explicitly and definitively linked to their highly resolved *Trypanosoma* identities prior to subsequent data synthesis and statistical evaluation in R [43] (Fig. 1).

#### 2.5.4. Data Matrix Construction and Visualization of Coinfection Profiles

To transition from discrete sample summaries to a unified analytical framework, the annotated alignment files (*.counts.modified.txt) were imported into the R statistical computing environment [43]. Data aggregation and matrix restructuring were executed utilizing the tidyverse package ecosystem [44]. An automated data-import pipeline iteratively read the individual sample files, extracting bat sample identifiers directly from the file nomenclature, and merged the discrete datasets into a comprehensive data frame. To ensure an accurate representation of parasite absence, the dataset was reshaped into a wide-format matrix, where missing data points— occurring when a specific *Trypanosoma* lineage was not detected within a given host—were programmatically imputed as absolute zeroes. Consistent with the primary objective of describing absolute coinfection frequencies, the underlying data matrix was intentionally not subjected to statistical normalization. Retaining the absolute, high-confidence mapped read counts allowed for an accurate representation of the raw magnitude of lineage dominance (Fig. 1).

To optimize visual clarity and focus on the most biologically relevant data, the dataset was stratified to isolate the top 10 most abundant *Trypanosoma* lineages across the entire cohort based on mean abundance. All remaining, less frequently detected lineages were computationally aggregated into a single holistic “Other” category.

## 3. Results

### 3.1. Impact of Alignment Stringency on the Detection of *Trypanosoma* Coinfections

Based on the three proposed scenarios, we observed a profound shift in the apparent complexity of intra-host *Trypanosoma* communities (Fig. 2).

The Lenient Scenario (High Apparent Diversity): Under relaxed filtering parameters (designed to accept multi-mapping reads and lower sequence identity), the resulting infection profiles exhibited massive apparent intra-host diversity (Fig. 2). The relative abundance revealed highly fragmented, multi-lineage distributions across nearly all sampled hosts. Individual bats appeared to be simultaneously co-infected by a wide array of distinct *Trypanosoma* lineages, closely mirroring the complex, hyper-diverse coinfection rates frequently reported in standard short-read metabarcoding literature. The absolute number of mapped reads retained in this scenario (i.e. the sequencing reads that were successfully mapped to a specific *Trypanosoma* 18S rRNA reference lineage within a single bat sample) was 14,153. The mean and standard deviation of taxa per sample in this scenario was 8.35 (1.4), with a maximum number of 10 coinfections. The most abundant lineage in this scenario was *Trypanosoma cruzi* isolate 262 (NCBI accession AY491763.1). The tenth most abundant lineage in this scenario was *Trypanosoma* sp. RNM063 (NCBI accession KT368796.1).

The Moderate Scenario (Intermediate Resolution): When standard intermediate alignment thresholds were applied, the complexity of the intra-host communities visibly decreased (Fig. 2). While some mixing of lineages remained evident, a substantial portion of the minor, overlapping read assignments was eliminated. Under these conditions, primary lineages began to clearly dominate the read counts within individual hosts, suggesting that much of the diversity observed in the lenient scenario was the result of bioinformatic noise and ambiguous read mapping among closely related sister taxa. The absolute number of mapped reads retained in this scenario was 2,668. The mean and standard deviation of taxa per sample in this scenario was 1.23 (0.58), with a maximum number of 3 coinfections. The most abundant lineage in this scenario was *Trypanosoma cruzi* isolate 262 (NCBI accession AY491763.1). The tenth most abundant lineage in this scenario was *Trypanosoma* sp. RM2054 (NCBI accession MF141862.1).

The Restrictive Scenario (Single-Lineage Dominance): The application of the highly restrictive baseline parameters fundamentally altered the observed infection landscape (Fig. 2). Once all expected methodological noise, truncated sequences, and ambiguous multi-mapping reads were computationally stripped away, the artifactual coinfections effectively collapsed. The resulting profiles demonstrated overwhelming single-lineage dominance (Fig. 2). In most of the sampled bats, a single *Trypanosoma* genotype accounted for approximately 100% of the high-confidence mapped reads, with overlapping secondary lineages either entirely absent or reduced to negligible frequencies. The absolute number of mapped reads retained in this scenario was 134. The mean and standard deviation of taxa per sample in this scenario was 1.09 (0.28), with a maximum number of 2 coinfections. The most abundant lineage in this scenario was *Trypanosoma cruzi* DTU I (isolate 262, NCBI accession AY491763.1). The least abundant lineage in this scenario was *Trypanosoma cruzi* DTU III (strain MT3663, NCBI accession AF288660.1).

### 3.2. Dominant *Trypanosoma* Lineages in the Sampled Cohort

Across the rigorously filtered dataset, the sequence-structure phylogeny and stringent mapping identified a limited number of distinct lineages actively circulating within the sampled host population. The most frequently detected dominant strains belonged primarily to the *Trypanosoma cruzi* and *Trypanosoma cruzi marinkellei* lineages, establishing exclusive infections in 21 and 10 of the samples, respectively (Fig. 2, outer circle). Only 3 coinfections (*T. cruzi* DTU I–*T. cruzi marinkellei*) were detected out of the 35 retained samples in this analysis (Fig. 2, outer circle).

A notable outcome of the conservative filtering scenario was the robust detection of *Trypanosoma cruzi marinkellei* within the armadillo cohort (Fig. 2, outer circle). While the majority of established infections demonstrated strict single-lineage dominance, *T. c. marinkellei* was unambiguously identified in these terrestrial hosts. Furthermore, the limited intra-host complexity observed in our finalized dataset included a rare mixed infection comprising *T. cruzi* DTU I and *T. c. marinkellei* within an armadillo sample (sample A57, Fig. 2, outer circle). The persistence of these specific alignments under the highly restrictive parameters (enforcing near-perfect sequence identity and an MAPQ ≥ 30) confirms these are genuine biological signals rather than artifactual mapping noise.

Additionally, the analysis isolated reads mapping with high confidence to *Trypanosoma cruzi* DTU III (TcIII) within the sampled armadillos (sample A12, Fig. 2, outer circle). This lineage maintained structural integrity and high mapping quality scores through all progressively stringent bioinformatic filters (Fig. 2, inner, middle and outer circles). The definitive assignment of TcIII in these hosts highlights the resolution of the nanopore 18S rRNA metabarcoding approach to accurately profile specific, naturally occurring *Trypanosoma* genotypes within the local mammalian fauna. Out of the 35 retained samples in this rigorous analysis (Fig. 2, outer circle), only 3 true coinfections were validated, underscoring the overwhelming trend toward competitive exclusion in the host bloodstream.

### 3.3. Summary of Infection Dynamics

The progressive elimination of pseudo-coinfections through strict bioinformatic filtering yields a clear biological signal. The data demonstrate that when high-resolution, full-length alignments are isolated, active multi-strain *Trypanosoma* coinfections within the bat bloodstream are exceedingly rare. The observed pattern of strict single-strain dominance is highly consistent with the ecological principle of competitive exclusion, suggesting that established *Trypanosoma* lineages actively prevent stable coexistence within the constrained niche of the host.

## 4. Discussion

The primary objective of this study was to evaluate the prevalence of *Trypanosoma* coinfections by modulating the bioinformatic stringency of nanopore-based 18S rRNA metabarcoding. Our results demonstrate that as alignment parameters become highly restrictive—enforcing near-perfect sequence identity and full-length coverage—the apparent hyper-diversity of intra-host *Trypanosoma* communities collapses. This indicates that much of the multi-lineage coinfection reported in previous metabarcoding literature may represent bioinformatic artifacts, such as ambiguous multi-mapping or chimeric sequences, rather than true biological coinfection. By distinguishing true ecological complexity—driven by asymmetric competition and niche restriction—from artificially inflated diversity metrics, our data strongly support Gause’s principle of competitive exclusion. Our assessment suggests that in the highly constrained niche of the mammalian bloodstream, established *Trypanosoma* lineages actively suppress competitors, resulting in the overwhelming single-lineage dominance observed in our conservative scenario. A remarkable finding in our rigorously filtered dataset is the detection of *Trypanosoma cruzi marinkellei* in armadillos, including a rare true mixed infection. Historically, *T. c. marinkellei* has been considered a bat-restricted subspecies [45,46]. The anomalous presence of a bat-associated lineage in a terrestrial host naturally raises the question of potential laboratory contamination. However, the DNA of both sets of samples were extracted in different dates: the armadillos in January 2024 and the bats in March 2024. In addition, the architecture of our conservative bioinformatic pipeline makes cross-sample contamination highly improbable. The detection of these specific reads was sustained even after enforcing a Mapping Quality (MAPQ) threshold of ≥ 30 (guaranteeing ∼99.9% unique mapping confidence) and requiring ≥ 98% sequence identity. If this were the result of simple amplicon pooling leakage, barcode-bleeding, or transient aerosol contamination in the laboratory, the spurious reads would likely be fragmented, uniformly distributed across samples, or eliminated by the stringent mismatch penalties (-s 80 -B 6). Instead, the recovery of high-confidence, full-length alignments suggests a genuine, albeit rare, ecological spillover event, highlighting the resolution power of long-read sequencing in capturing the true extent of host-parasite networking.

Furthermore, the detection of *T. cruzi* DTU III in an armadillo within the Ecuadorian Amazon represents a highly significant biogeographical record [47,48]. While the sylvatic transmission cycles in Ecuador and northern South America are overwhelmingly dominated by DTU I [49], DTU III is classically associated with the terrestrial ecotope and armadillo hosts (Order Cingulata) much further south, particularly within the Gran Chaco and Southern Cone regions [47,48]. Identifying DTU III in an Ecuadorian armadillo extends the known geographic distribution of this lineage. More importantly, it reinforces the hypothesis of a deep, highly conserved evolutionary association between armadillos and DTU III that persists independent of broad geographical boundaries or the dominant circulating lineages in the local sylvatic environment.

### 4.1. Potential limitations of the study

Despite the robustness of our analytical pipeline, certain methodological and biological limitations must be acknowledged. Primarily, the implementation of highly stringent bioinformatic filters—specifically enforcing an MAPQ ≥ 30 and a sequence identity threshold of ≥ 98%—functions as a double-edged sword. Just as defining specific distance cut-offs in cluster analyses is critical to aligning taxonomic assignments with actual ecological observations, our conservative thresholds were essential to successfully strip away analytical noise, multi-mapping ambiguity, and chimeric sequences. However, this rigorous approach inherently risks producing false negatives by computationally discarding genuine, low-abundance secondary lineages. If a legitimate coinfecting strain is present at marginal parasitemia levels, or if residual long-read sequencing errors push its alignment identity marginally below the 98% cutoff, it is entirely excluded from the final matrix. Consequently, while our pipeline confidently eliminates artifactual pseudo-coinfections, the reported absolute single-lineage dominance might represent a slight underestimation of the true micro-diversity within the host.

Furthermore, our reliance on a ∼560 bp target amplicon of the 18S rRNA gene presents inherent resolution constraints. While this highly conserved marker is a reliable standard for resolving broad trypanosome taxonomy and inter-lineage diversity, it may lack the hyper-variable resolution required to consistently separate extremely closely related intra-DTU variants. Therefore, while our data strongly support inter-lineage competitive exclusion, highly homologous strains may still evade detection.

Finally, our findings reflect a localized temporal snapshot of infection dynamics. The analyzed bat blood and armadillo tissues capture parasitemia at a single point in time, inherently missing the broader temporal fluctuations characteristic of *Trypanosoma* infections. It is biologically plausible that sequential dominance, temporal succession, or specific tissue tropisms allow for broader multi-lineage persistence over the lifespan of the host. Even if asymmetric competition actively limits their simultaneous co-occurrence in the bloodstream at any given moment, a more complex whole-host infection history cannot be entirely ruled out. Another potential source of limitation for the specificity of *Trypanosoma* detection is the tempo and mode of tissue conservation for analysis, such as the age of the collected tissue and its medium of preservation, both parameters often highly heterogeneous in opportunistic collections.

### 4.2. Final remarks

In conclusion, while 18S rRNA metabarcoding using Oxford Nanopore Technologies provides exceptional resolution for wildlife parasitology, fine-tuning algorithmic parameters is critical to avoid overstating intra-host diversity. We propose that once methodological noise is stripped away, the resulting infection profiles reveal that active multi-strain *Trypanosoma* coinfections are ecologically infrequent. Mechanistically, our results could be explained by the host bloodstream favoring competitive exclusion, maintaining strict single-strain dominance while occasionally permitting the circulation of highly specific, rare spillover events that would otherwise be lost in the noise of lower-resolution pipelines.

## Acknowledgements

We extend due credit to Matus Valach for the use of the *Trypanosoma* silhouette image under the CC0 1.0 Universal Public Domain Dedication license in www.phylopic.org.

## 5. Supporting information

Reference Taxonomic Assignment Databases: The custom 18S rRNA *Trypanosoma* reference databases generated and utilized in this study are publicly available on Zenodo at https://zenodo.org/records/20721872. S1 Data: this repository includes the initial highly-redundant dataset comprising 2,097 sequences (trypanosoma_18S_db_2097.fas); S2 Data: the dereplicated dataset of 336 unique sequences (trypanosoma_18S_db_336.fas); and S3 Data: the final manually curated database of 113 reference sequences used for mapping (trypanosoma_18S_db_113.fas).

S4 Text. Sensitivity Analysis of Alignment Thresholds. This supplementary document provides the mathematical justification for selecting the 98% sequence identity threshold applied in the conservative bioinformatic scenario. It details the comparative evaluation of read retention and multi-mapping behavior across 95%, 97%, 98%, and 99% cut-offs, demonstrating why 98% serves as the optimal inflection point to eliminate analytical noise without discarding genuine biological sequences due to minor sequencing errors. The S4 text is available at https://doi.org/10.5281/zenodo.20721872

Bioinformatics Code and Pipelines: The custom Python/awk scripts (including process_data.py) and the exact code for the three structured minimap2 filtering paradigms (Lenient, Moderate, and Conservative scenarios) are hosted on GitHub and can be accessed at: https://github.com/PJV-Ecu/Fine-tuning-parameters-to-identify-mixed-infections-of-Trypanosoma

Raw Sequence Data: The raw Oxford Nanopore sequencing reads generated in this study have been deposited in the Sequence Read Archive (SRA)] and can be accessed under Bioproject PRJNA1502415.

